# Integrated Autolysis, DNA Hydrolysis and Precipitation Enables an Improved Bioprocess for Q-Griffithsin, a Broad-Spectrum Antiviral and Clinical-Stage anti-COVID-19 Candidate

**DOI:** 10.1101/2021.12.30.474602

**Authors:** John S. Decker, Romel Menacho-Melgar, Michael D. Lynch

## Abstract

Across the biomanufacturing industry, innovations are needed to improve efficiency and flexibility, especially in the face of challenges such as the COVID-19 pandemic. Here we report an improved bioprocess for Q-Griffithsin, a broad-spectrum antiviral currently in clinical trials for COVID-19. Q-Griffithsin is produced at high titer in *E. coli* and purified to anticipated clinical grade without conventional chromatography or the need for any fixed downstream equipment. The process is thus both low-cost and highly flexible, facilitating low sales prices and agile modifications of production capacity, two key features for pandemic response. The simplicity of this process is enabled by a novel unit operation that integrates cellular autolysis, autohydrolysis of nucleic acids, and contaminant precipitation, giving essentially complete removal of host cell DNA as well as reducing host cell proteins and endotoxin by 3.6 and 2.4 log_10_ units, respectively. This unit operation can be performed rapidly and in the fermentation vessel, such that Q-GRFT is obtained with 100% yield and >99.9% purity immediately after fermentation and requires only a flow-through membrane chromatography step for further contaminant removal. Using this operation or variations of it may enable improved bioprocesses for a range of other high-value proteins in *E. coli*.

**Highlights:**

- Integrating autolysis, DNA hydrolysis and precipitation enables process simplification
- Autolysis reduces endotoxin release and burden to purification
- Q-Griffithsin recovered from fermentation vessel at >99.9% purity and 100% yield
- Q-Griffithsin purified to anticipated clinical grade without conventional chromatography
- The resulting bioprocess is 100% disposables-compatible, scalable, and low-cost

## Introduction

The need for innovation in biopharmaceutical manufacturing has been highlighted by the COVID-19 pandemic [1], as well as, more generally, by the long-term increase in the use and cost of biologic drugs [2]. This has been recognized through regulatory initiatives such as the Advanced Technology Team within FDA [3] and the formation of government-academic-industrial partnerships such as the National Institute for Innovation in Manufacturing Biopharmaceuticals (NIIMBL) [4]. At the level of process design, the key trends are intensification and flexibility, where intensification refers to increasing efficiency on the basis of cost, space, and/or time [5], and flexibility refers to processes that are modular, portable from facility to facility, and amenable to changes in production rates [4]. At the level of technology, some approaches are especially interesting because of their potential to deliver both flexibility and intensification simultaneously. These include single-use systems [4,6]; alternatives to conventional chromatography that are less equipment- and buffer-intensive [7–10]; and the integration of multiple unit operations into more streamlined procedures with smaller footprints, e.g., by combining primary recovery and initial purification using expanded bed adsorption of cell lysates [11] or host cells engineered for selective product release [12].

These trends and challenges in biopharmaceutical manufacturing have a direct impact on the global response to COVID-19. Though there is a recognized need for globally-available therapeutics to complement vaccines [13], and especially for broad-spectrum antivirals [14], current biotherapeutics for COVID-19 are not broad-spectrum and have not been widely available worldwide, in part because of the cost of manufacturing and the difficulty of building new manufacturing capacity for unpredictable demand (i.e., process inflexibility) [13]. One drug candidate that offers both broad-spectrum activity and manufacturing advantages over existing biotherapeutics is Q-Griffithsin (Q-GRFT), an oxidation-resistant single-residue variant of Griffithsin (GRFT) [15]. GRFT and Q-GRFT are antiviral lectins with remarkably broad-spectrum and potent activity as well as promising safety profiles [16]. Importantly, a Q-GRFT-based nasal spray is currently in clinical trials for the prevention of COVID-19 [17]. Previously, we have reported that cost and scale limitations of the existing GRFT bioprocess represent a significant barrier to the feasibility of GRFT as a widely-used antiviral, and reported a novel intensified *E. coli* bioprocess that overcomes those limitations [18]. However, that process still required costly fixed equipment including multiple chromatography columns, making its flexibility limited. Because therapeutics for viral pandemics necessarily have highly unpredictable demand both temporally and geographically, and because global distribution of biotherapeutics can be a significant challenge, it is important that manufacturing approaches be as simple and flexible as possible to enable rapid and local changes in capacity.

Here, we report a new process for Q-GRFT that improves upon the previous process in both intensification and flexibility. The fundamental advance of this process is the use of a novel integrated procedure for cellular autolysis, autohydrolysis of nucleic acids, and contaminant precipitation, so that Q-GRFT is obtained at high purity immediately after fermentation. The bioprocess does not require conventional chromatography and is entirely compatible with single-use systems after fermentation. Relative to the previous process, it is more modular, easily transferable, and responsive to changes in volume demand, while also having a lower cost of goods sold (COGS) at scales up to several kL of fermentation capacity.

## Materials and methods

### Reagents and media

FGM30 media was prepared as previously described [19]. Kanamycin sulfate was used at a working concentration of 35 ug/mL. Unless otherwise stated, all materials and reagents were of the highest possible grade and purchased from Sigma (St. Louis, MO).

### Strains and plasmids

E. coli strains DLF_R003, DLF_R004 [20], and DLF_Z0025 [21], as well as plasmid pHCKan-yibD-QGRFT plasmid (Addgene #158748) [18], were prepared as previously described.

### Fermentation

Instrumented fermentations were performed as previously described [19].

### Autolysis and precipitation

A protocol for autolysis and auto DNA hydrolysis triggered by heat shock was developed on the basis of a previously-reported method for autolysis triggered by freeze-thaw [20]. Briefly, a thermostable GFP variant [22] was cloned into DLF_R004 and expressed in shake flasks according to previously-published methods [19]. Cells were then incubated at various temperatures for 1 hr with or without Triton X-100 at 0.1% v/v and fluorescence in clarified supernatants was compared to that in samples lysed by sonication to estimate the relative degree of protein release.

Following fermentation, cell lysis and initial Q-GRFT purification were performed in the fermenter in an integrated fashion by combining the newly-developed heat-triggered autolysis method with a previously-reported method for GRFT purification by precipitation [18]. Specifically, the fermenter temperature and pH were ramped to 60 °C and 3.4, respectively, and maintained there for 1 hr. Next, ammonium sulfate was added to a final concentration of approximately 815 mM, pH was maintained at 4, and the suspension was again held for 1 hr. Finally, lysates were decanted for clarification. Gentle agitation was maintained throughout using hollow-bladed impellers at 300 rpm.

### Clarification, tangential flow filtration and membrane chromatography

Following autolysis and precipitation, Q-GRFT-containing lysates were clarified by centrifugation in a swinging-bucket rotor at 4000 RCF for 15 minutes. The supernatant was then filtered using a 0.2 µm PES filter (Genesee Scientific). After clarification, Q-GRFT was diafiltered against 10 diavolumes of one of several buffers using a tangential flow filtration (TFF) capsule with a PES membrane of 10 kDa MWCO value (Minimate, Pall Corporation, Port Washington, NY). TFF was performed with a feed flow rate of approximately 10 L m^2^ min^-1^ and a transmembrane pressure of approximately 20 psi. Finally, following TFF, pure Q-GRFT was prepared by flow-through membrane chromatography on a salt-tolerant anion exchange capsule (Sartobind STIC PA nano, Sartorius, Goettingen, Germany) according to manufacturer instructions.

### Quantification of Q-GRFT and contaminants

In samples determined to be homogeneous by SDS-PAGE, Q-GRFT was quantified by BCA assay (Pierce BCA Protein Assay Kit, Thermo Fisher Scientific, Waltham, MA). In crude cell lysates, Q-GRFT was quantified by an in-house ELISA using standard sandwich ELISA techniques. Briefly, HIV gp120 (NIH HIV Reagent Program, Cat. #13354) was immobilized at 1 µg/mL on 96-well assay plates (Nunc Maxisorp, Thermo Fisher Scientific, Waltham, MA). Plates were then blocked with 5% w/v BSA and incubated with clarified cell lysates containing 200 ng/mL total protein followed by incubation with a rabbit polyclonal anti-GRFT antibody (gift of Barry O’Keefe, National Cancer Institute, Bethesda, MD) at 1:400 dilution. Bound Q-GRFT was detected using an HRP-conjugated goat anti-rabbit polyclonal antibody at 1:5000 dilution and TMB (Cat. #65-6120 and #N301, respectively, Thermo Fisher Scientific, Waltham, MA).

Endotoxin was quantified using Endosafe cartridges (Cat. #PTS20005F, Charles River Laboratories, Wilmington, MA) according to manufacturer instructions. Residual host-cell proteins in Q-GRFT samples were quantified using an ELISA kit (Cat. #F410, Cygnus Technologies, Southport, NC). Residual *E. coli* chromosomal DNA in purified Q-GRFT samples was quantified using qPCR as previously described [23]. Briefly, a primer-probe pair targeting *E. coli* 16s rRNA was obtained by chemical synthesis (Integrated DNA Technologies, Coralville, IA). 20 µL qPCR reactions were carried out in duplicate according to the protocol provided with the Luna Universal Probe qPCR Master Mix (New England Biolabs, Ipswich, MA). Quantification was performed by comparison to a standard curve of *E. coli* genomic DNA (Cat. #AAJ14380MA, Fisher Scientific, Waltham, MA) at concentrations from 1 to 10000 pg per reaction. Residual host cell nuclease was quantified using a fluorogenic nuclease detection kit (DNAseAlert, Integrated DNA Technologies, Coralville, IA) and a benzonase standard. 50 μL reactions were prepared in triplicate in a clear-bottom black 384-well plate, with each well containing 35 μL of nuclease-free water, 5 μL of DNAseAlert substrate, 5 μL of 10X DNAseAlert buffer, and 5 μL of nuclease-free water (negative control), sample, or benzonase standard ranging from 1.56 to 0.049 milliunits per reaction in a 2-fold dilution series. Plates were incubated for 1 hr at 37 °C before reading with a HEX filter set.

### Technoeconomic analysis and bioprocess modeling

Bioprocess models were prepared using SuperPro Designer (Intelligen, Inc, Scotch Plains, NJ) according to methods previously described [18]. Q-GRFT expression in both the new model and the previously-described model was set to 0.1 g/gDCW based on the expression observed in the present study, and downstream equipment in the previously-described model was resized to accommodate the increased mass of Q-GRFT. Because large-scale pricing for the STIC membrane was not available, we used SuperPro’s pricing models for Sartobind Q (Sartorius, Goettingen, Germany). Membrane sizing was based on an endotoxin capacity of 4000000 EU/mL, according to information from the manufacturer, and single-use operation was assumed.

## Results

### Q-GRFT is expressed to high titer in two-stage fermentations

Previously, GRFT and Q-GRFT have been expressed in *N. benthamiana* and *E. coli*, with the highest reported titer being 2.7 g GRFT/L in two-stage medium-density fermentations of *E. coli* strain DLF_Z0025 [18]. First, we sought to determine whether the recently-developed *E. coli* strain DLF_R004 [20], which enables cellular autolysis and autohydrolysis of DNA and RNA, could support similar titers of Q-GRFT. In microfermentations (Fig. 1A), the relative expression levels of Q-GRFT and GRFT did not vary substantially among DLF_R004 and two non-autolysis controls including DLF_Z0025, though there was a small but significant increase in Q-GRFT expression for DLF_R003 compared to DLF_R004. However, Q-GRFT expression was consistently somewhat lower than GRFT expression in all strains. Nonetheless, scaling up to 1 L fermentations (Fig. 1B) with DLF_R004 resulted in a Q-GRFT titer 36% higher than that previously obtained for GRFT in DLF_Z0025 (i.e., 3.68 ± 0.5 g Q-GRFT/L vs. 2.7 g GRFT/L).

**Figure 1.**
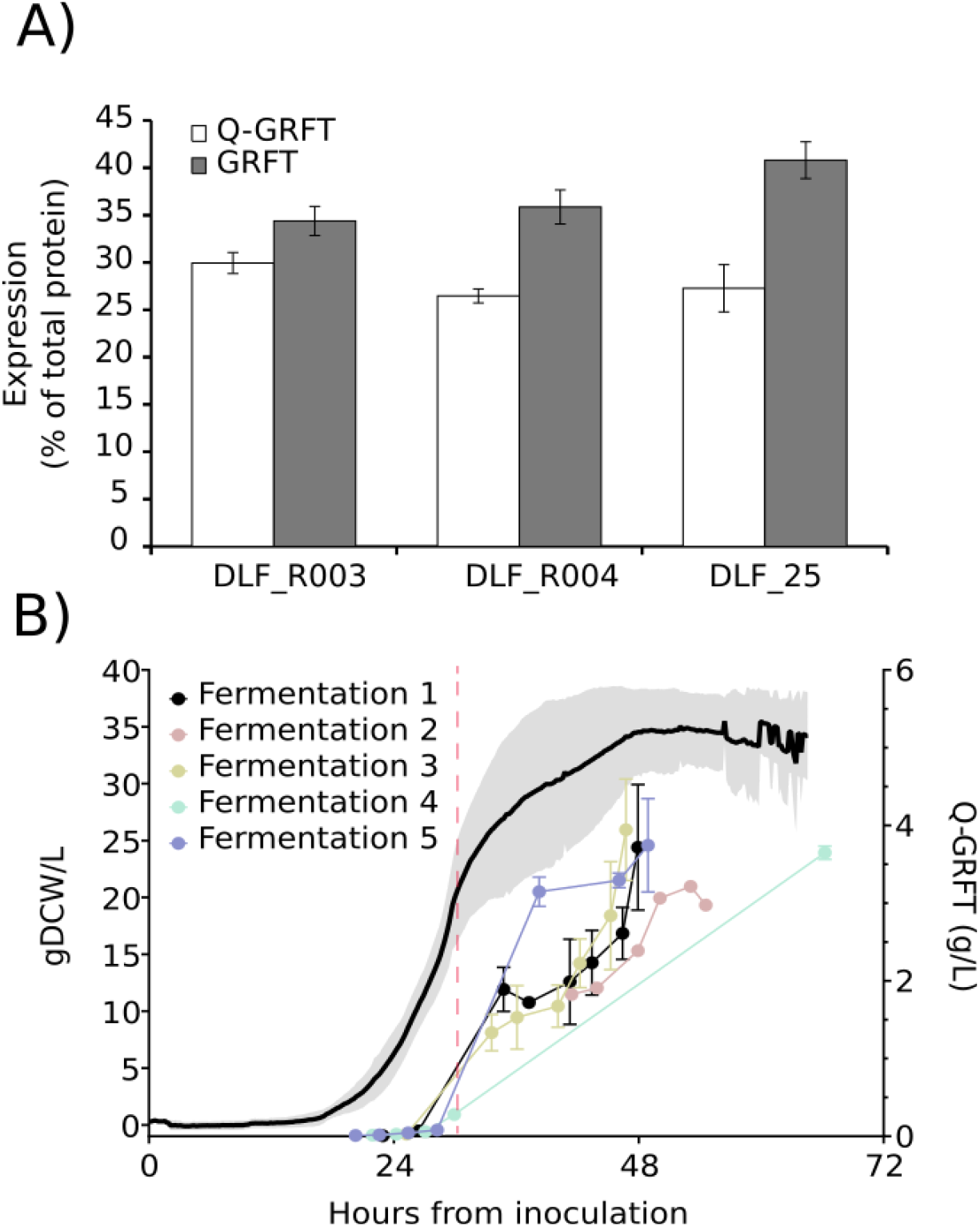
High-titer expression of Q-GRFT by two-stage fermentation. A) Relative expression of Q-GRFT and GRFT was compared by SDS-PAGE densitometry in microfermentations of an autolysis *E. coli* strain (DLF_R004) vs. two non-autolysis controls (DLF_R003 and DLF_25). N = 3 for each group. B) Q-GRFT and biomass accumulation were monitored during instrumented 1 L fermentations of DLF_R004. Solid line and shaded area: gDCW/L, mean ± SD, N = 6. Circles: Q-GRFT g/L, mean ± SD, N = 1-3 per timepoint. Red dashed line: mid-exponential growth.

### Autolysis selectively reduces endotoxin release and can be triggered by heat shock

Having expressed Q-GRFT to high titer, we turned to the establishment of an efficient product release strategy. We reasoned that autolysis using DLF_R004 might cause less micronization of the cell membrane than mechanical disruption for a given level of protein release, and therefore that endotoxin might remain associated with larger membrane particles and be more easily removed during clarification. In DLF_R004 lysed either by sonication or by autolysis using the freeze-thaw based method previously reported [20], we found that autolysis reduced endotoxin levels per unit total soluble protein by approximately 7.4-fold (Fig. 2A). Next, we desired to determine whether the existing autolysis method could be adapted to better fit a large-scale and low-cost bioprocess. Specifically, we sought to avoid its reliance on a freeze-thaw cycle, which is not amenable to large-scale processing, and on Triton X-100, which is no longer generally allowed by European drug regulators [24]. We reasoned that the freeze-thaw cycle could be avoided if elevated temperatures caused a similar membrane disruption without deactivating the lysozyme and nuclease of DLF_R004. To test this, we expressed a heat-stable GFP variant (muGFP [22]) in DLF_R004 and monitored extracellular fluorescence as a function of temperature and of the presence or absence of 0.1% Triton X-100 (Fig. 2A). At 60 °C, protein release was not significantly different from that obtained with sonication, even in the absence of Triton. Expression of muGFP in the non-autolysis control strain DLF_R003 confirmed that protein release at 60 °C was still dependent on the lysozyme activity of DLF_R004 (Fig. 2B).

**Figure 2.**
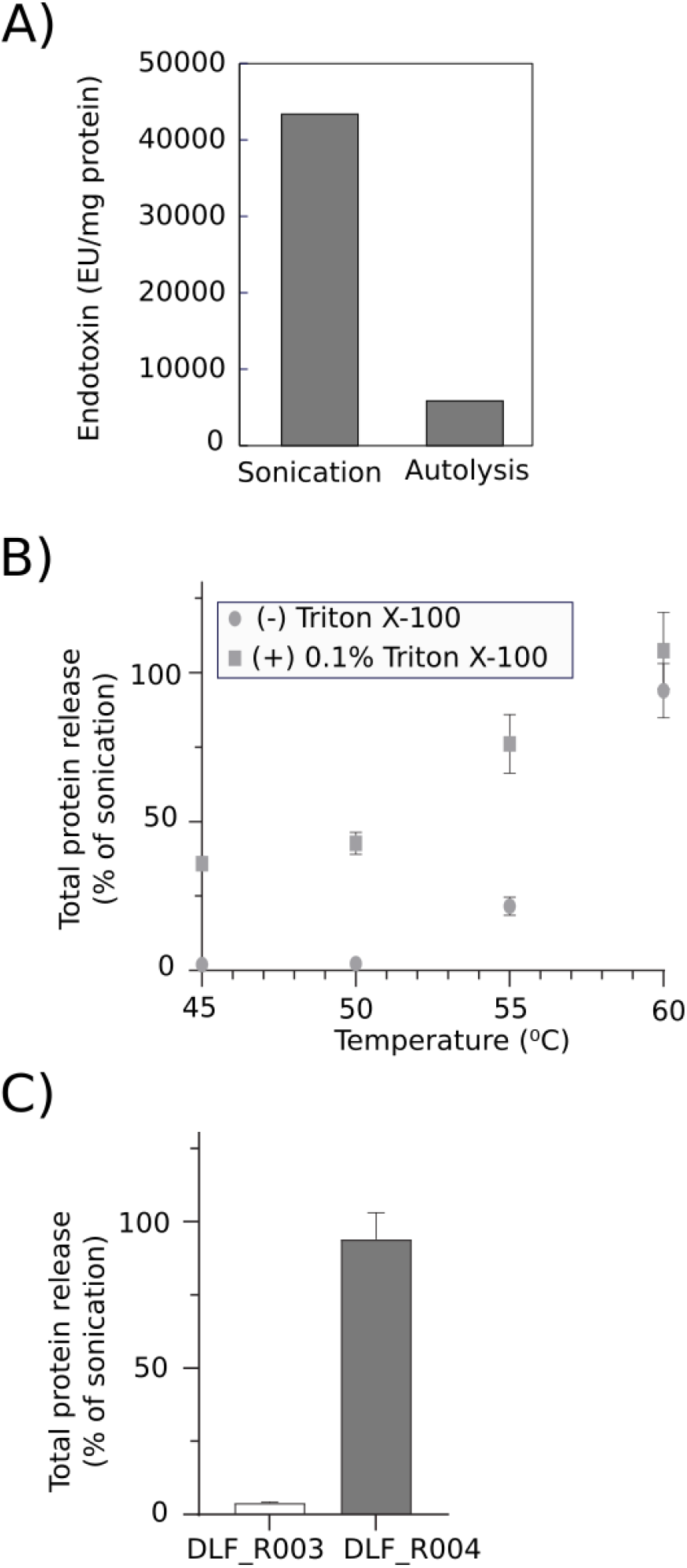
Characterization of autolysis effects on endotoxin release and development of a heat shock-triggered autolysis method. A) EU per mg total soluble protein following lysis of DLF_R004 by either sonication or autolysis using the freeze-thaw method [20]. B) muGFP fluorescence relative to sonication-lysed controls for DLF_R004 with or without 0.1% Triton X-100 at various temperatures. N = 3. C) muGFP fluorescence relative to sonication-lysed controls for DLF_R004 vs. DLF_R003 at 60 °C in the absence of Triton X-100. N = 3.

### Q-GRFT is rapidly released and purified by in-fermenter autolysis and precipitation

Next, we sought to integrate the heat shock-triggered and detergent-free autolysis method developed above with a previously-reported method for rapid purification of GRFT by precipitation of contaminants [18]. We reasoned that, since both the new autolysis method and the precipitation method occur at the same temperature (60 °C), the two might be combined in a single unit operation. Furthermore, since nothing is required for either method but pH and temperature control, mixing, and addition of a salt solution, we reasoned that such an integrated operation could be performed immediately after fermentation inside the bioreactor. Thus, Q-GRFT could be rapidly and effectively released and purified at the earliest possible stage of downstream processing, reducing the load on subsequent operations and thereby contributing to a more efficient and lower-cost process. The accumulation of Q-GRFT throughout a 1 L fermentation, and the purified Q-GRFT obtained after performing the integrated autolysis and precipitation operation within the fermenter, are shown in Fig. 3. Soluble protein harvested from the fermenter 2.5 hours after the end of fermentation contained no detectable host-cell proteins (HCPs) by SYPRO Ruby staining (Thermo Fisher Scientific, Waltham, MA) (Fig. 3A).

**Figure 3.**
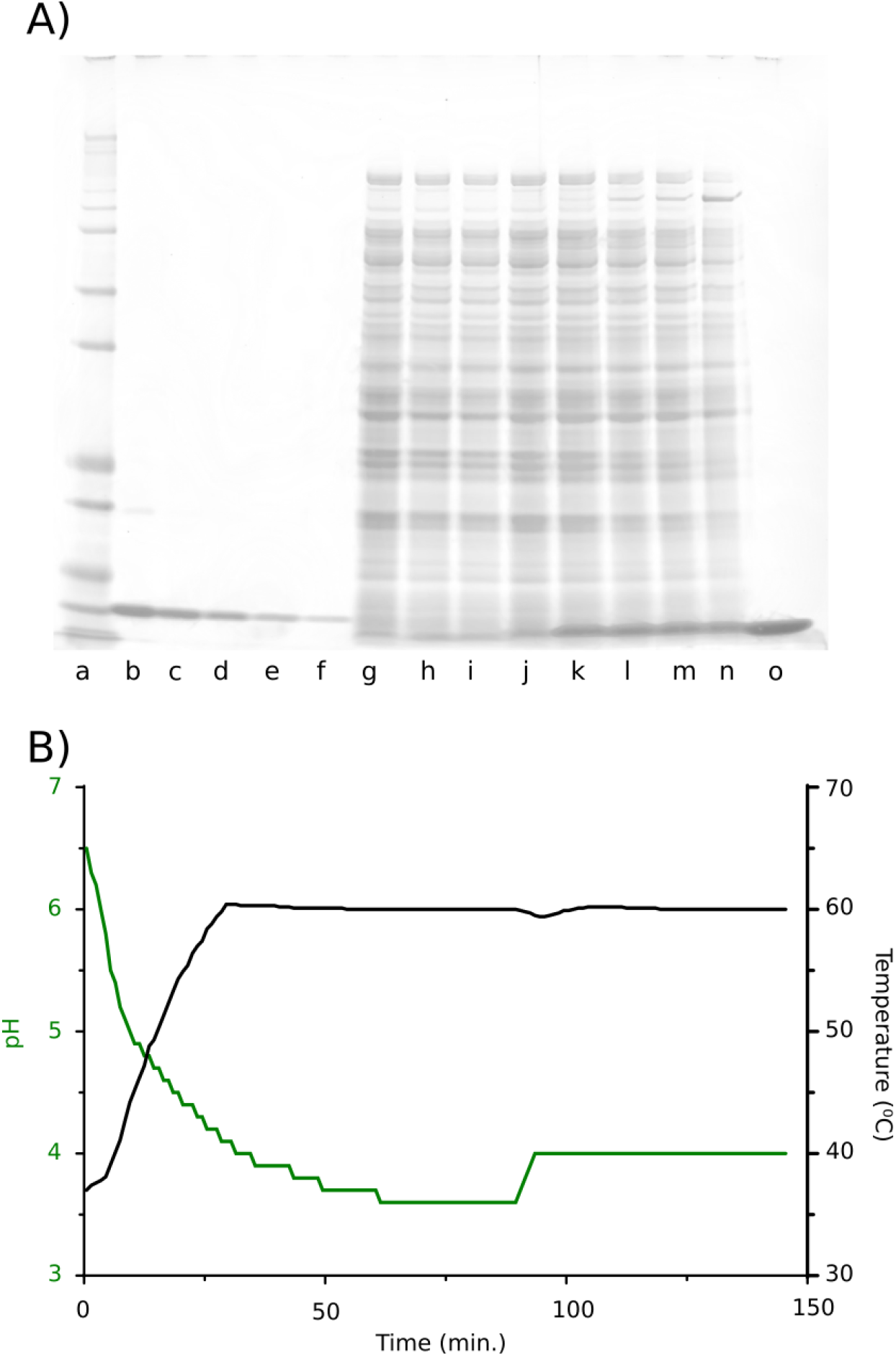
In-fermenter autolysis and precipitation for rapid release and purification of Q-GRFT. A) SDS-PAGE showing accumulation of Q-GRFT throughout a 1 L fermentation and purified Q-GRFT harvested from the fermenter after autolysis and precipitation. Lanes: a = Mk 12 unstained standard (Thermo Fisher Scientific, Waltham, MA); b-f = His-GRFT standard, 1 μg to 62.5 ng in 2-fold series; g-n, 1.5 μg total protein from fermentation samples at 20, 22, 25, 28, 38, 46, 49, and 64 hrs after inoculation; o, 1 μg total soluble protein following autolysis and precipitation (67 hrs after inoculation). B) Temperature and pH profile inside the fermenter during the autolysis and precipitation operation. The upward inflection of pH at approximately 90 mins. marks the addition of ammonium sulfate.

### Final purification and quality assessment of Q-GRFT

Having expressed Q-GRFT at high titer and developed a rapid, simple, and highly effective method for releasing and purifying it immediately after fermentation, it was next necessary to complete the new bioprocess by quantifying and removing additional key contaminants (i.e., endotoxin and residual host-cell DNA (rcDNA)) to below anticipated regulatory limits. Q-GRFT in current clinical trials is being administered as a nasal spray. Therefore, we assumed a dose of <20 mg, based on a typical nasal spray delivery volume of 100 μL per spray [25] and a reasonable stable Q-GRFT concentration of approximately 20 mg/mL (personal communication, Barry O’Keefe, National Cancer Institute). Further, we assumed the following impurity targets: <1000 ppm HCPs, based on a lower than usual immunogenicity risk given that preclinical data indicates that nasally-delivered Q-GRFT is not systemically absorbed (personal communication, Barry O’Keefe, National Cancer Institute); <0.5 ppm DNA (i.e., <10 ng/dose [26]); and <4.8 ppm endotoxin (i.e., <48 EU/mg Q-GRFT given the commonly used conversion of 10 EU per ng endotoxin), based on a suggested limit for topical drugs of 100 EU/m^2^ [27] and a nasal epithelial surface area of 9.6 m^2^ [28].

After precipitation, we determined the levels of endotoxin and HCPs to be 628.33 and 945.75 ppm, respectively, representing approximately 2.4 and 3.6 log reductions from the initial cellular composition. Notably, HCP clearance in this process was improved by 10-fold compared to when the same precipitation method was used following sonication rather than autolysis (2.6 log reduction [18]); however, it remains to be determined whether this improvement was due to the lysis method or to other factors such as different mixing conditions. Q-GRFT concentration and rcDNA were not quantitatively determined at this stage due to matrix interference, though agarose gel electrophoresis showed no detectable DNA (not shown). Further, mass spectroscopy suggested that remaining contaminants were almost entirely of approximately 2.5 kDa molecular weight (not shown). We reasoned that a TFF stage with 10 kDa cutoff would retain Q-GRFT while simultaneously removing small molecular weight contaminants and allowing conditioning of the sample for final purification. TFF diafiltration was performed with 10 diavolumes of a buffer containing 10 mM Bis-Tris, pH 6.0, and 484 mM NaCl. Following TFF, Q-GRFT yield was determined to be 105.44 ± 6.07 % relative to the fermentation titer, while endotoxin and HCP levels were determined to be, respectively, 180.14 ± 63.58 and 602.86 ± 130.90 ppm (0.54 and 0.20 log reductions compared to after precipitation).

Given that the predominant remaining contaminant after TFF was endotoxin, which has a strong negative charge at pH greater than approximately 2.5, we chose to use flow-through anion exchange chromatography as the final purification operation. Further, we selected a membrane rather than a packed-bed resin for this operation because of its advantages of flexible scaling, single-use compatibility, and reduced buffer consumption. Using the Sartobind STIC PA membrane (Sartorius, Goettingen, Germany), we found that endotoxin was cleared to well below target levels using a pH of 6.0, slightly above Q-GRFT’s predicted isoelectric point of 5.4, and an NaCl concentration of 484 mM (conductivity of approximately 40 mS/cm). Initial tests at pH 4.0 and 20 mS/cm, chosen to minimize Q-GRFT binding to the membrane and according to manufacturer recommendations, respectively, failed to reduce endotoxin below 5 ppm (not shown), possibly due to charge interactions between Q-GRFT and endotoxin. After the STIC flow-through step, Q-GRFT yield was determined to be 85.24 ± 9.56 % relative to the fermentation titer. Endotoxin and HCP levels were, respectively, 0.51 ± 0.36 ppm and 534.57 ± 97.81 ppm. The level of rcDNA was less than an upper bound value of 0.13 ± 0.02 ppm, based on no detectable amplification in a qPCR assay sensitive to 0.02 pg/μL.

The Q-GRFT yield and the levels of endotoxin, HCPs, and DNA after each process stage are summarized in Figure 4 for six replicate fermentations and three replicate runs of the downstream operations.

**Figure 4.**
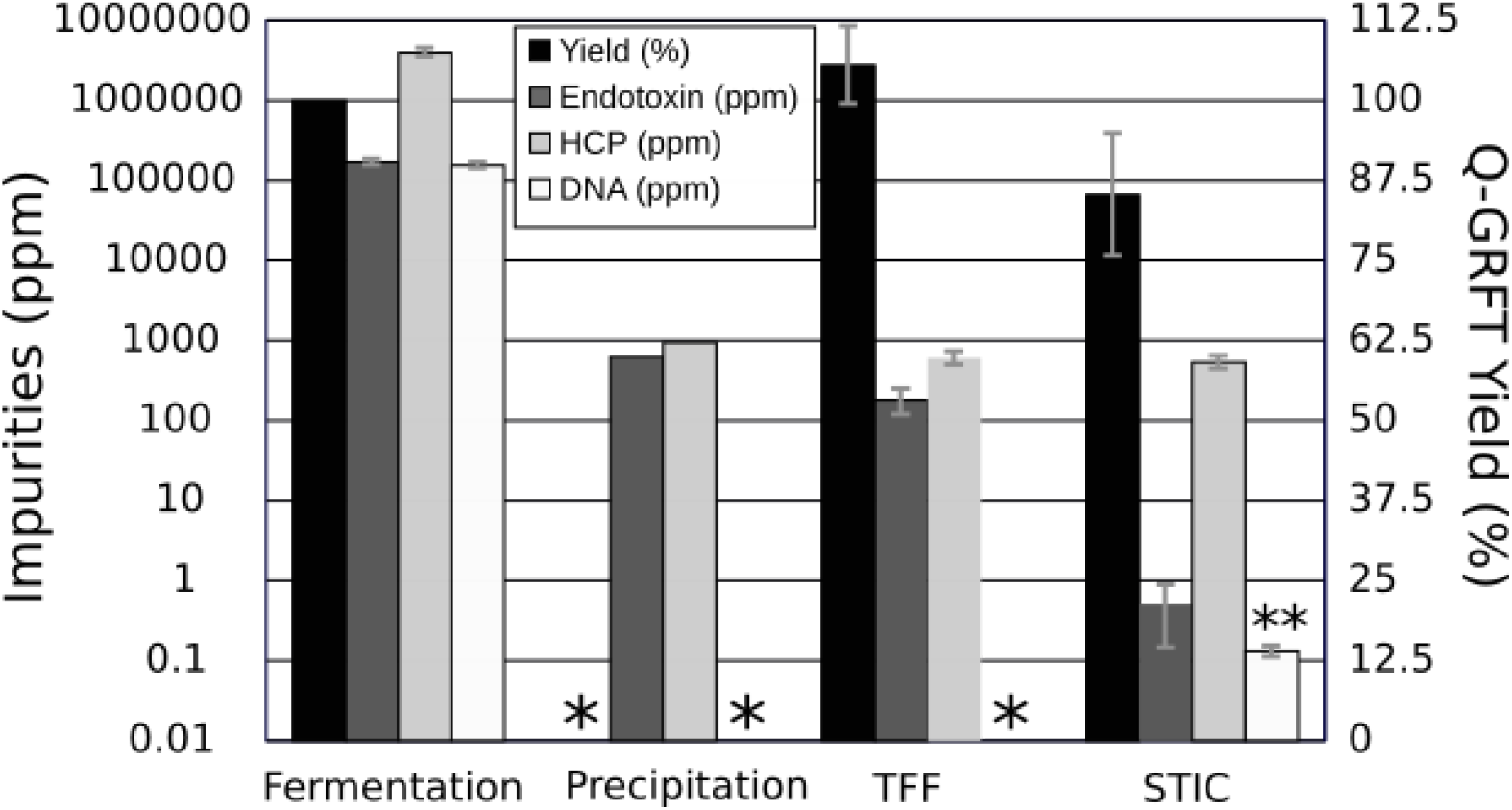
Yield of Q-GRFT and levels of key impurities after each stage of the bioprocess. At the fermentation stage, impurity levels are estimates based on the literature [29] and the titer of Q-GRFT as determined by ELISA. At other process stages, impurity and Q-GRFT abundances were determined by the assays described in Materials and Methods. *: indicates that the corresponding value was not determined at a given process stage. **: indicates that the value shown is an upper bound (DNA was not detectable by qPCR in the STIC-purified samples, LOQ = 0.02 pg/μL). Columns show the grand mean of 3 bioprocess replicates (except for fermentation, N = 6 bioprocess replicates) with technical replicates of N = 3, 4, 3, or 2 for yield, endotoxin, HCP and DNA measurements, respectively. Error bars show SD of the 3 bioprocess replicates. Yield at the fermentation stage is 100% by definition.

### Overview of the improved Q-GRFT bioprocess

In summary, we have developed a bioprocess capable of producing Q-GRFT at anticipated clinical grade with high yield and only a few rapid, simple downstream operations. Notably, the process does not require conventional chromatography and enables an entirely single-use downstream train. This process is summarized schematically in Figure 5. Fill and finish operations included in the SuperPro models and economic evaluations are not shown.

**Figure 5.**
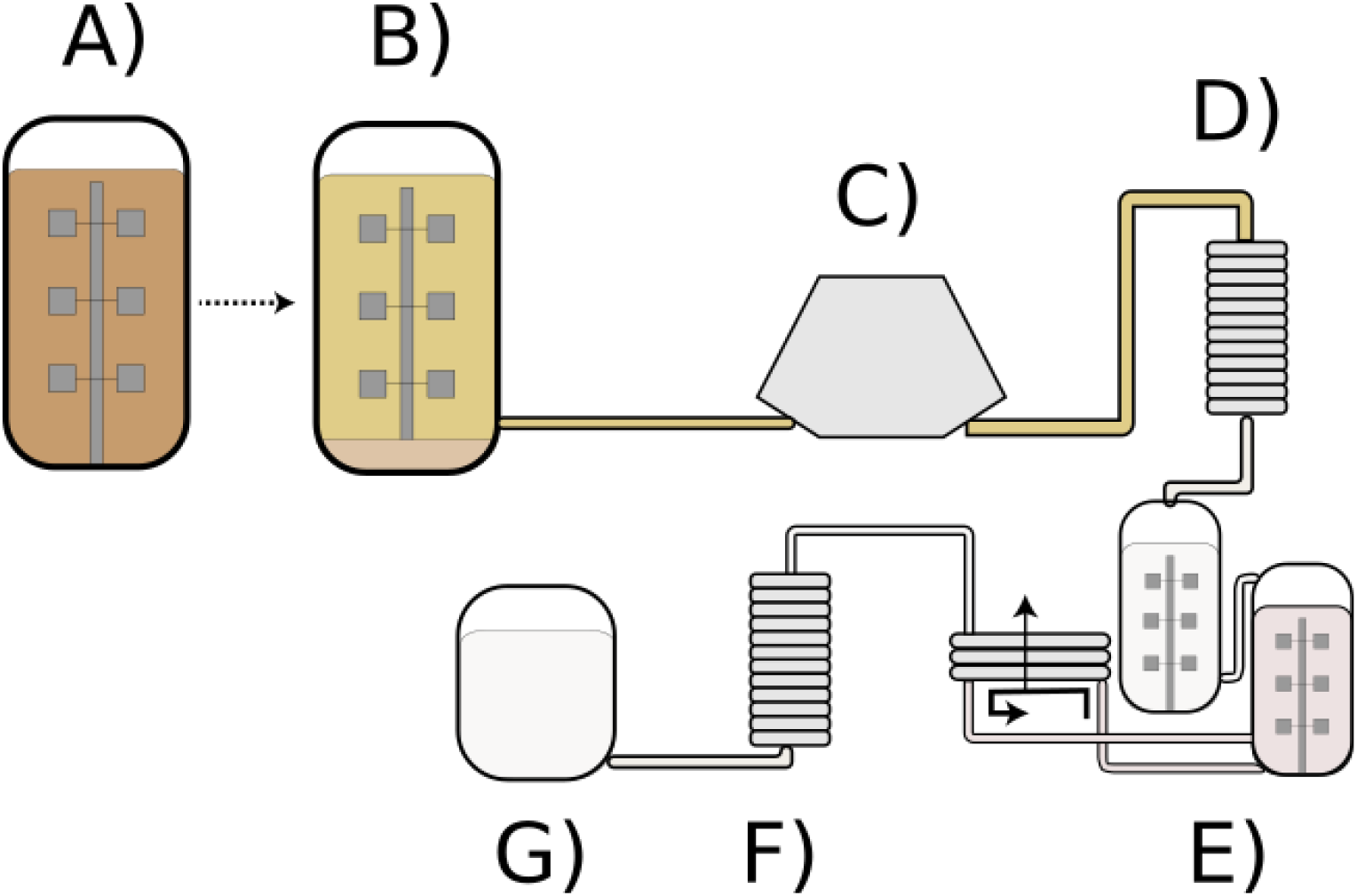
Overview of the improved Q-GRFT bioprocess. A) Fermentation. B) An integrated procedure for cellular autolysis, auto-hydrolysis of DNA and RNA, and contaminant precipitation, which can be performed within the fermenter. C) and D) Lysate clarification by centrifugation and/or dead-end filtration. E) Buffer exchange by tangential flow filtration. F) Flow-through membrane chromatography. G) Bulk drug substance.

### Economic aspects of the improved Q-GRFT bioprocess

To assess the economic competitiveness and scalability of the improved Q-GRFT bioprocess reported here, we created a detailed model of a cGMP plant employing the process using SuperPro Designer (Intelligen, Scotch Plains, NJ). For comparison, we also adapted our previously-published model of a scalable and low-cost GRFT bioprocess [18] by updating it to reflect the level of Q-GRFT expression achieved in the present study. The previously reported process made use of the same precipitation method as used here, but without autolysis or autohydrolysis of nucleic acids, and also required two conventional chromatography columns instead of the single membrane chromatography stage used here. In general, the two processes have comparable COGS at all scales (Fig. 6A), although the new process has somewhat higher COGS at very large scales (e.g., 30 kL and 120 kL) and moderately lower COGS at small to medium scales (by 27% and 23% at 300 L and 3 kL, respectively). A more detailed breakdown of COGS by process section and cost category (Fig. 6B) revealed that the COGS advantage of the new process at small scales (300 L and 3 kL) comes not only from eliminating primary recovery costs (i.e., cell harvest and homogenization), but also from roughly 50% reductions in labor, facility and materials-related expenses in purification.

**Figure 6.**
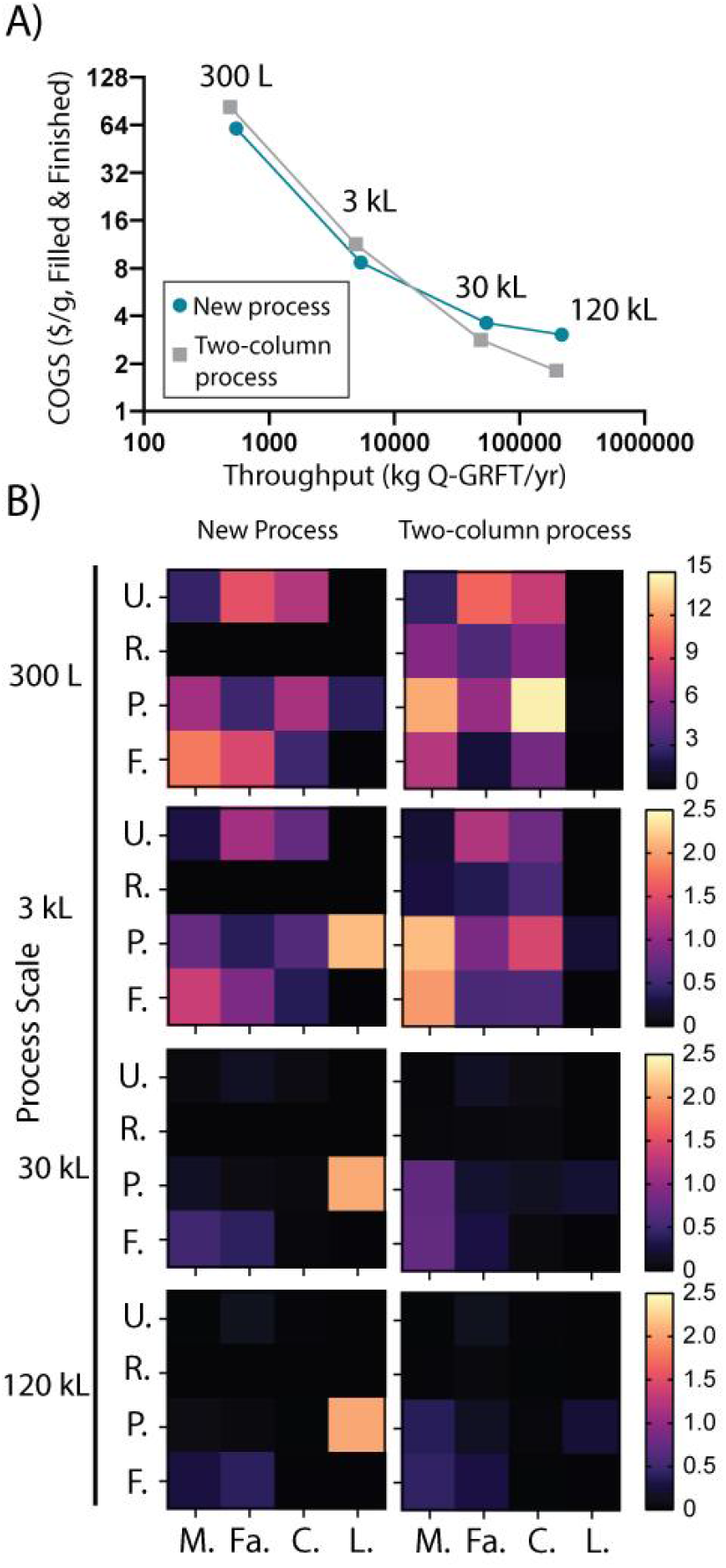
Economic overview of the improved Q-GRFT bioprocess vs. a process using two chromatography columns [18]. A) Total COGS and throughput for both processes at four scales. B) Cost structure for the processes and scales shown in A. U = upstream, R = primary recovery, P = purification, F = fill/finish. M = materials, Fa = facility, C = consumables, L = labor and QC. Color bar = COGS ($/g).

At large scales (30 kL and 120 kL), however, COGS in both processes become essentially driven by one factor: by the cost of the chromatography membrane per unit Q-GRFT for the new process; and by raw materials usage per unit Q-GRFT, e.g. for chromatography buffers, in the previous process (Fig. 6B).

## Discussion

### A flexible and efficient bioprocess to support global deployment of a broad-spectrum antiviral and anti-COVID-19 drug candidate

The biopharmaceutical manufacturing industry is increasingly emphasizing process efficiency (i.e., intensification) and flexibility in order to contain biologic drug costs, enable more agile responses to emerging and unpredictable needs, and make biologics economically viable for a wider range of markets in terms of both geography and disease indication. Global availability of effective biologic therapeutics and preventives is a key unmet need in the ongoing response to COVID-19 and preparation for future pandemics, but will require addressing these industry-wide manufacturing challenges. Here, we have reported a novel, intensified and flexible bioprocess for a clinical-stage anti-COVID-19 candidate and broad-spectrum antiviral, Q-GRFT. This process is efficient with respect to cost, labor, equipment and raw materials; recovers Q-GRFT at high yield; and achieves anticipated clinical-grade purity with only a single purification step after the product leaves the fermentation vessel. Perhaps most importantly, all downstream operations are amenable to single-use processing, so that it is as simple as possible to modify the process scale or transfer the process to a new facility. These factors place the present work in contrast with previous methods for the production of GRFT and Q-GRFT [18,30], as well as with processes commonly in use for other biotherapeutics such as antibodies, which are reliant on costly fixed equipment and do not combine low COGS, scalability, and a potential for rapid and simple changes in capacity. As such, we believe the bioprocess reported here represents a unique and significant advance towards ensuring that Q-GRFT can be effectively deployed against COVID-19 and/or future pandemics, pending its clinical validation.

### Integration of autolysis, autohydrolysis of DNA, and contaminant precipitation enables process simplification

The key enabling feature of our improved Q-GRFT bioprocess is the integration of cellular autolysis, autohydrolysis of DNA and RNA, and contaminant precipitation into a single unit operation that essentially eliminates rcDNA as well as greatly reducing levels of endotoxin and HCPs. If any one of these three contaminants had not been so efficiently cleared at this early stage of the process, it is likely that subsequent downstream operations would have been required in greater number and/or with greater complexity and reliance on fixed equipment, such as bind-and-elute column chromatography.

The integrated operation described here, though developed for Q-GRFT, may enable improved bioprocesses for other products produced in *E. coli*. In particular, we expect that the process could be applied directly for any other proteins that are solubly expressed in *E. coli*, thermostable to around 60 °C, and of roughly similar size to Q-GRFT (approximately 26 kDa), given that heat denaturation and protein surface area-dependent entropic effects [31] are expected to drive precipitation in our process. Perhaps the most promising molecules to fit this description are the single-domain antibodies (“nanobodies”), an emerging class with significant potential in both therapy [32] and diagnosis [33], which are routinely expressed in *E. coli* [34], are approximately 15 kDa in weight, and have an average melting temperature of 65 - 70 °C [35]. Another high-value candidate is granulocyte colony stimulating factor (G-CSF), which has been expressed to high titer in soluble form in the *E. coli* periplasm [36], is approximately 22 kDa in weight, and has a melting temperature of 57 °C [37]. In the case of G-CSF, we expect that the operation could be conducted at slightly lower temperatures than used here provided that 0.1% Triton X-100 was used in autolysis, given that approximately 75% protein release was obtained at 55 °C under these conditions. Furthermore, the autolysis and precipitation operation reported here might be sufficient as the sole purification step for a number of valuable non-pharmaceutical products, which have less stringent purity requirements. Examples include enzymes for research or industrial uses, biomaterials, or food ingredients.

### Further improvements to the autolysis and precipitation process

Though the present work offers methods that might be readily and usefully applied to a number of high-value *E. coli* products besides Q-GRFT, there are several apparent avenues for extending its applicability and/or improving its efficiency. First, additional host strain or product engineering might enable autolysis to directly reduce HCP levels. For example, it may be possible to modify the autolysis method used here to give more limited cellular disruption, and/or specifically to release the contents of the periplasm but not the cytoplasm. Further host strain engineering might also enable autolysis with improved reductions in endotoxin levels. Additionally, alternative precipitation strategies could be considered. Though the current precipitation method was optimized for HCP reduction and Q-GRFT yield, it was not designed specifically for the reduction of endotoxins, another key contaminant. It also requires temperature and pH conditions that would denature many proteins of interest. However, in addition to the elevated temperatures and kosmotropic salts as used here, protein precipitation can also be caused by many other agents that work through different mechanisms, such as polyelectrolytes, organic solvents, and nonionic polymers [31]. Phase separations of endotoxin can also be performed using some detergents [42] or polymers [43]. By further exploring the design space including some of these additional factors, precipitation conditions may be found that give clearance of HCPs and endotoxin comparable to or better than the current procedure, while using less denaturing conditions.

## Acknowledgments

This study was supported by NIH grant 3R61AI140485-02S1. We gratefully acknowledge the collaboration of scientists at the National Cancer Institute, the National Center for Advancing Translational Sciences, and the Biopharmaceutical Development Program of the Frederick National Laboratory.

## Author Contributions

JSD and MDL conceived the study and prepared the manuscript. JSD, RMM and MDL performed experiments. JSD performed bioprocess modeling. All authors edited and revised the manuscript.

